# w*TSA-CRAFT* : an open-access web server for rapid analysis of thermal shift assay experiments

**DOI:** 10.1101/2023.06.22.546086

**Authors:** Victor Reys, Julien Kowalewski, Muriel Gelin, Corinne Lionne

**Affiliations:** Centre de Biologie Structurale (CBS), CNRS UMR 5048, Université de Montpellier, INSERM U 1054, 29 rue de Navacelles, Montpellier, 34090, France

## Abstract

The automated data processing provided by the *TSA-CRAFT* tool enables now to reach high throughput speed analysis of thermal shift assays. While the software is powerful and freely available, it still requires installation process and command line efforts that could be discouraging. To simplify the procedure, we decided to make it available and easy to use by implementing it with a graphical interface via a web server, enabling a cross-platform usage from any web browsers. We developed a web server embedded version of the *TSA-CRAFT* tool, enabling a user-friendly graphical interface for formatting and submission of the input file and visualization of the selected thermal denaturation profiles.

We describe a typical case study of buffer condition optimization of the biologically relevant APH(3’)-IIb bacterial protein in a 96 deep-well thermal shift analysis screening. w*TSA-CRAFT* is freely accessible for non-commercial usage at https://bioserv.cbs.cnrs.fr/TSA_CRAFT.

## Introduction

Characterizing protein stability is often central in a wide variety of biochemistry applications, including buffer optimization, protein crystallisation, ligand screening and fragment-based drug design for drug discovery process (1-2). Among the various methods available, the thermal shift assay (TSA) allows to analyse protein melting temperature by unfolding it upon temperature variations from differential scanning fluorimetry (3). Its increasing usage, due to the potential high-throughput capabilities, was still limited to the scope of analysing qRT-PCR output data as it requires substantial efforts and curve fitting knowledge. To compensate this issue, a tool *TSA-CRAFT*, for TSA -Curve Rapid and Automatic Fitting Tool (4), was developed for the analysis of such output data. This tool is used for automated data processing and Boltzmann equation fitting of qRT-PCR result files, enabling a quicker analysis of TSA experiments out of 96 (or 384) wells plates. Although the tool is open source, cross platform, freely accessible and based on few shell and Perl scripts, the library dependencies, its installation and usage can still be discouraging, especially for users unfamiliar with terminal command lines.

For this reason, we decided to spare users from the installation processes and made a graphical interface for input file submission by embedding the software under a dedicated web server hosted in our laboratory. In addition, we developed a new tool to format the raw qRT-PCR output data into a valid input file for *TSA-CRAFT*, saving even more time and limiting the risk of errors. In this application note, we describe a free and accessible web server of the embedded *TSA-CRAFT* tool plus the formatting tool under the w*TSA-CRAFT* name, available at https://bioserv.cbs.cnrs.fr/TSA_CRAFT.

### Format specification

Input files needed to run w*TSA-CRAFT* are similar to the one(s) required by *TSA-CRAFT* (highlighted in green in Supplementary Fig. S1). It consists of a mandatory thermal shift analysis table formatted as comma separated value (csv), where the first column holds the temperatures at which data points were measured and all subsequent columns contain fluorescence measurements for each well.

An additional optional csv file can also be provided, enabling the annotation of the various wells allowing to disambiguate their content while displayed. This file must contain only three columns (separated by commas); *i)* the index for the wells (#Well_index), *ii)* the human annotated condition (#Annotation) and *iii)* the name for the well(s) serving as reference(s) (#Reference_well_index), separated by spaces if multiple of them are present in the experiment.

Finally, a second optional input enabling the overlaying of multiple wells (up to a maximum of eight) on the same graph.Wells to be combined must be separated by commas and also present in the analysis.

In addition to the standard input section required to run *TSA-CRAFT*, weadded a supplementary input section (highlighted in pink at bottom right of home page in Supplementary Fig. S1). It automatically converts raw thermal shift output data into a valid *TSA-CRAFT* input csv file, which allows to reach one step further in the speed of usage. This format converter is dedicated to the transformation of qRT-PCR Mx3005P (Stratagene) output data and takes as input two text files exported from the experiment results, the dissociation and amplification curve files.

### Implementation

We implemented the *TSA-CRAFT* tool on a virtual machine hosted in our laboratory and running under Ubuntu LTS 16.04 using Apache2 (v.2.4.29) server and few Python3 (v.3.6.8) cgi scripts to validate input files, launch the *TSA-CRAFT* Perl (v.5.26.1) scripts and post-process html rendering. This allows a simplified tool usage protocol in a cross-platform manner from major web browsers (tested on Firefox, Edge, Chrome and Safari). w*TSA-CRAFT* is freely accessible for non-commercial usage at https://bioserv.cbs.cnrs.fr/TSA_CRAFT.

#### Graphical interface

Information about the original *TSA-CRAFT* tool (4) are available, with access to the publication, scripts repository and user manual (highlighted in blue in Supplementary Fig. S1). A simple yet effective web interface allows a graphical rendering for input files submission on the home page. Next to each input sections, test files are provided for download. Finally, as the standard *TSA-CRAFT* output are already provided in html language, a small post-processing of the result page is performed: local paths are modified to URLs, a supplementary download links is added in the header and an additional input section for later graph overlaying generation is made accessible.

#### Download of the results

As we did not want to block users on the web server results, we are providing a zip archive of all the generated results by *TSA-CRAFT*. This archive contains all previously generated results (html result page with relative paths and related graph overlays) and therefore can be used in a local environment. We strongly suggest users to download their results after their analyses as we are cleaning old queries results from our server on a weekly basis, for obvious saving of hard drive storage space.

#### Graph overlay plots

*TSA-CRAFT* provides an extra script (*plot_multi_well_curves*.*pl*) generating comparative graphs overlay for a set of 2 to 8 wells. To take advantage of all options allowed by the tool, we decided to provide this supplementary option at multiple locations. First, immediately in the home page input section (green in Supplementary Fig. S1, as describe above), and also in the *TSA-CRAFT* result page. In the result page (Supplementary Fig. S2), we added a new input form, from which users are able to use again this function to generate unlimited multi-well graphs, without recomputing the overall results. Links to download the various graphs produced are stacked at the bottom of the result page (Supplementary Fig. S2). Upon graph generation, the content of the zip archive is also updated with the newly made one.

Finally, we observed that wells annotated with more than 20 characters were going out of the graph. To compensate this issue a small computation is performed, proportionally lowering the size of characters (up to a minimum of size 4) in the legend of the figures, hence allowing longer description of annotated wells.

#### Raw data format converting tool

We are providing an additional format converting tool, enabling the automated generation of a valid *TSA-CRAFT* input csv file from raw qRT-PCR output data. To do so, first the dissociation file is parsed and cycles names are mapped to the corresponding temperatures. Then, the amplification file, which is holding fluorimetric information is parsed to gather amplitudes at each cycle. In order to be sure that temperatures correspond to the cycles, raw data should be exported without smoothing. Finally, we write a new csv file containing temperature values in the first column, followed by fluorescence amplitudes obtained for each well present in the experiment. We return the corresponding file as a downloadable csv file to the user, who simply needs to place it in the *TSA-CRAFT* input section. Thus, this new tool allows a faster and more secure conversion of raw data into an input file for *TSA-CRAFT*.

### Case study

We applied a standard protocol for the screening by thermal shift assay of buffer conditions optimization for the protein APH(3’)-IIb (UniProtKB_Id: Q9HWR2_PSEAE) of *Pseudomonas aeruginosa* implicated in bacterial resistance to aminoglycoside antibiotics (5-6). We used the commercially available RUBIC BufferScreen MD1-96 kit from Molecular Dimensions (3) which is presented as a 96 × 1 mL deep-well block. We have slightly adapted the protocol suggested by the manufacturer to fit the 20 μL tips of our Opentrons pipetting robot (Fig. 1A). We dispensed 4 μL of a 2× mixture containing in house purified APH(3’)-IIb (2.5 μM final) and SyproOrange (5× final), then added and mixed 20 μL of RUBIC BufferScreen. The thermal denaturation screening was performed in a qRT-PCR device (Mx3005P, Stratagene) using a Cy3 and SYAL emission filter at a gain of 8 and 1 respectively (Fig. 1B). The fluorescence amplitudes obtained using the SYAL filter being saturated, those obtained with the Cy3 filter were used for further analysis.

**Figure 1.**
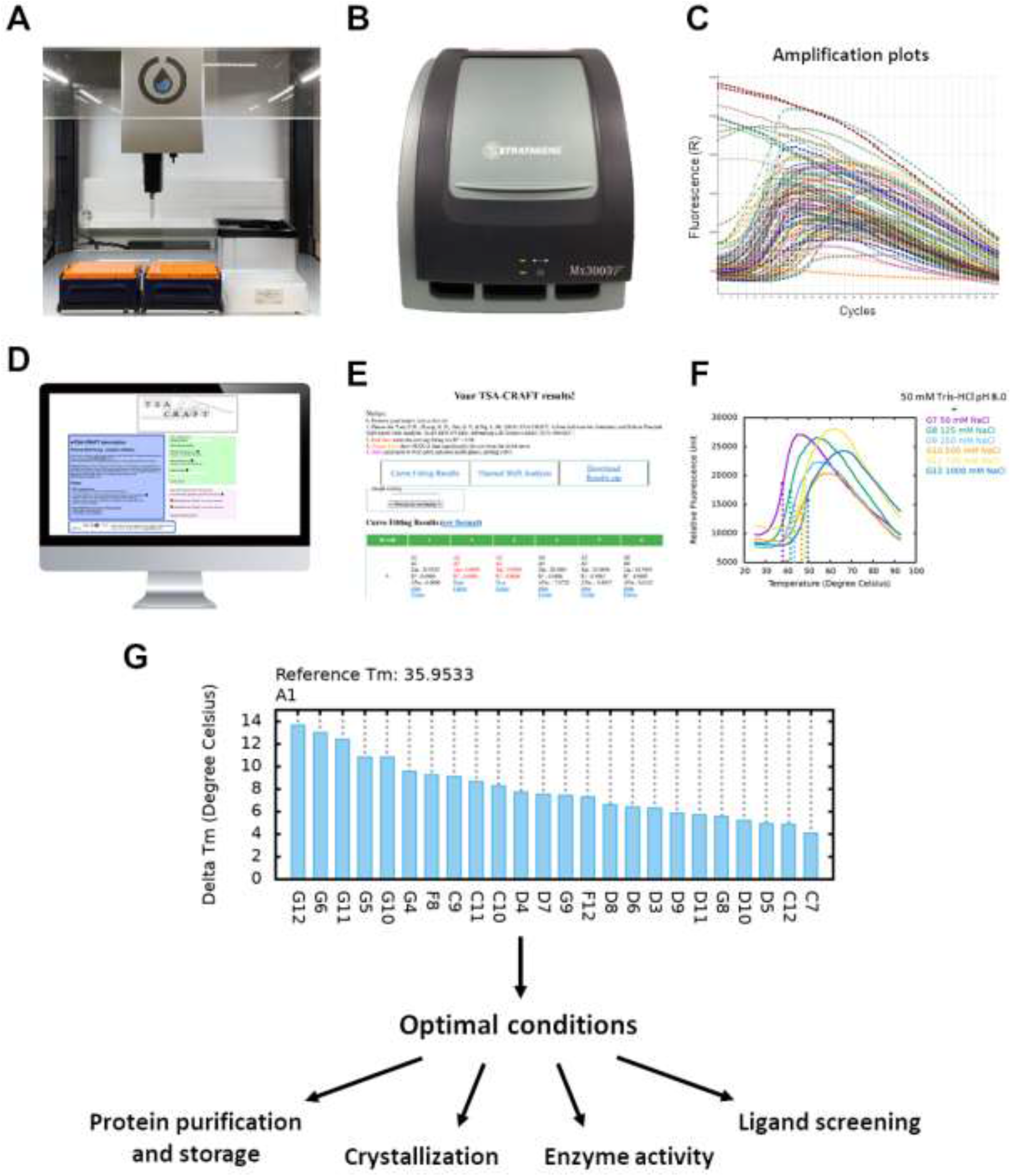
Workflow for the optimization of buffer conditions by TSA screening. (**A**) In a 96-well PCR plate, the protein is mixed with SyproOrange and different buffers using a pipetting robot. (**B**) The PCR plate is placed in a qRT-PCR device and warmed up to 95°C while the fluorescence is recorded. (**C**) The amplification and dissociation data are exported as text file in MxPro software. (**D**) The exported files are converted into a valid input csv file sent to w*TSA-CRAFT* (at https://bioserv.cbs.cnrs.fr/TSA_CRAFT). Curve fit analysis is performed by the embedded *TSA-CRAFT* software. (**E**) The results are used to compare the thermal stability of the protein in different buffer conditions. (**F**) Up to 8 selected plots can be overlaid. (**G**) Optimal buffers are chosen accordingly to the requirements of the techniques used such as protein purification and storage, crystallization for X-ray structure determination, *in vitro* enzyme activity or ligand screening

Raw dissociation and amplification outputs files (Fig. 1C, Supplementary File S1) were exported in text file format (Format 2 - Horizontally grouped plot). Both were submitted to w*TSA-CRAFT* for automated file conversion into valid *TSA-CRAFT* input file (Fig. 1D). The returned file and the optional annotation csv file were then placed in the TSA-Analysis form and analysed by *TSA-CRAFT* (Fig. 1D). Within seconds, the 96 curve fittings were performed and the post-processed result page was sent back (Fig. 1E). Interestingly, we could rapidly identify the influence of NaCl concentration, allowing to reach higher melting temperatures (Tm) with high concentrations (Fig. 1F, Supplementary Fig. S3A and S3D). Eventually, in well G12 (1 M NaCl, 0.05 M Tris-HCl at pH 8.0) the highest Tm of 49.6°C was reached, which corresponds to a ΔTm of 13.7°C with respect to water (well A1). In addition, pH lower than 5.5 were identified as deleterious for the protein thermal stability (Supplementary Fig. S4A and S4B).

These results allowed us to carry out several follow up works, by choosing the most suitable buffer for each techniques taking into account the preferences of the studied protein. These new experiments (Fig. 1G), including protein purification and storage, crystallization, structure determination by X-ray crystallography, activity and affinity measurements and ligand screening, will be described in another paper.

## Supporting information

Amplification file

Annotation file

Dissociation file

TSA-CRAFT input file

## Acknowledgements

The authors thank Violaine Moreau and Jean-Luc Pons for hardware survey and continuous support. We also thank Gilles Labesse and all beta-testers for their helpful remarks. Finally, we thank Po-Hsien Lee and associated authors of the original *TSA-CRAFT* (4) software for letting us the opportunity to embed their tool under a free and accessible cross-platform web service.

## Funding

Victor Reys and Julien Kowalewski were supported by la Ligue Nationale contre le Cancer, the Agence Nationale de la Recherche [ANR-17-CE18-0011], the Association Vaincre la mucoviscidose and the Association Grégory Lemarchal. The CBS is part of the ChemBioFrance research infrastructure and is a member of the France-BioImaging (FBI) and the French Infrastructure for Integrated Structural Biology (FRISBI), 2 national infrastructures supported by the French National Research Agency (ANR-10-INBS-0004 and ANR-10-INBS-0005, respectively).

## Conflict of interest

We declare no conflict as the embedding of TSACRAFT software respects the stated agreement terms, agreed by Po-Hsien Lee in May 2020 (7).

---

**Figure S1.**
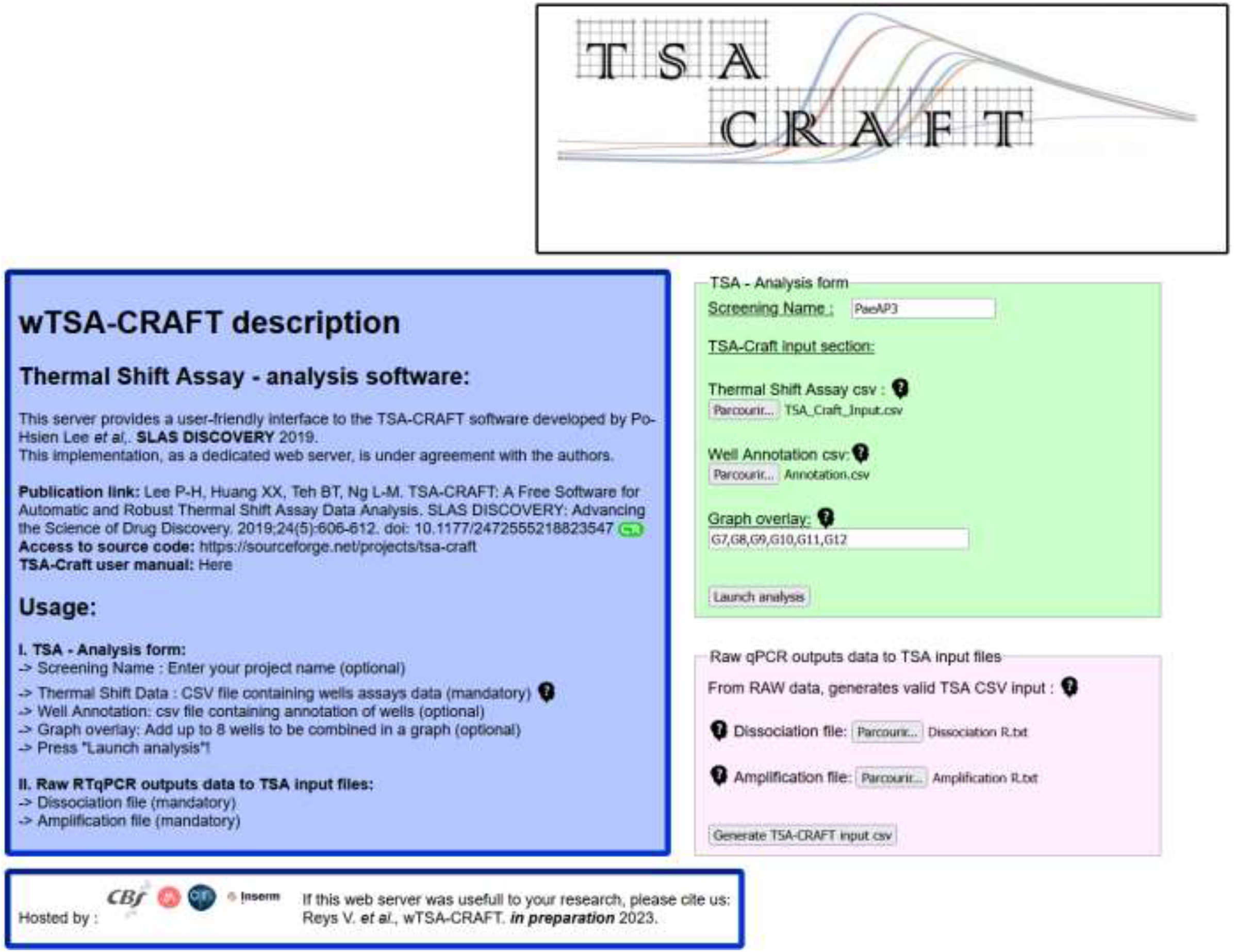
Screenshot of the w*TSA-CRAFT* home page at *https://bioserv.cbs.cnrs.fr/TSA_CRAFT/*. The different sections of the home page are highlighted by frames of different colours. On the left, in blue, information of the original *TSA-CRAFT* algorithm and access to the publication, software and user manual. On the top-right, in green, the TSA Analysis form, containing a text input to provide the name of the screening, enabling to properly name the later generated zip archive (optional), the mandatory TSA input file as csv, the well annotation csv (optional) and a text input for the combination of multiple graph overlay (optional). At the bottom-right, in pink, the form holding the two input file acceptor for the automated format conversion from raw qRT-PCR outputs to valid *TSA-CRAFT* input csv file by providing both the dissociation and amplification files.

**Figure S2.**
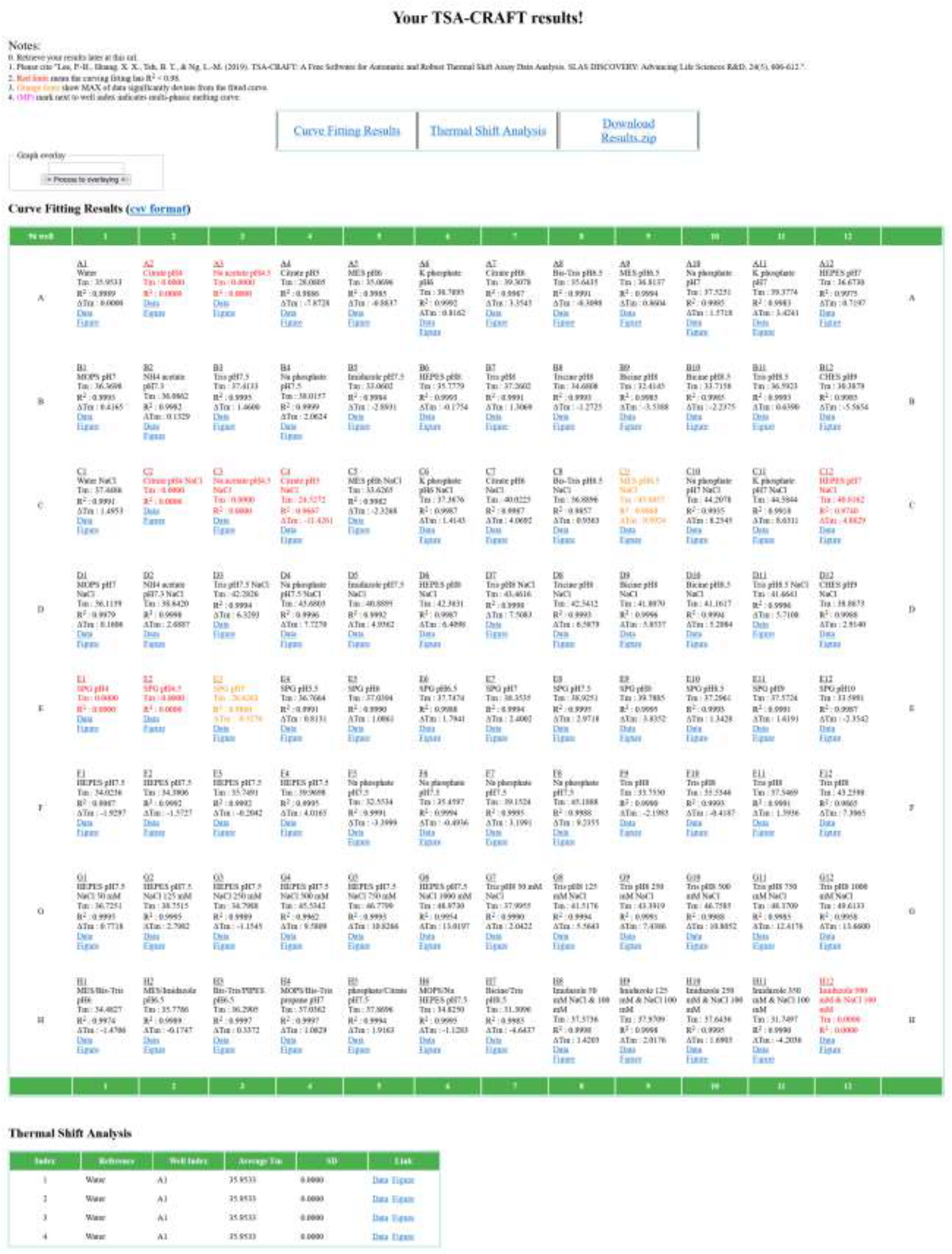
Screenshot of the edited *TSA-CRAFT* result page. In the header are available the various data files that can be downloaded. An additional link to a zip archive containing all results has been added by editing the original *TSA-CRAFT* output. A supplementary ‘Graph overlay’ combination input form is also provided, where the user can launch the ‘*plot_multi_well_curves.pl*’ script by providing up to 8 wells separated by comas. The result page is followed by a table holding the various thermal shift analysis for each well with a colour code related to the quality of the curve fit.

**Figure S3.**
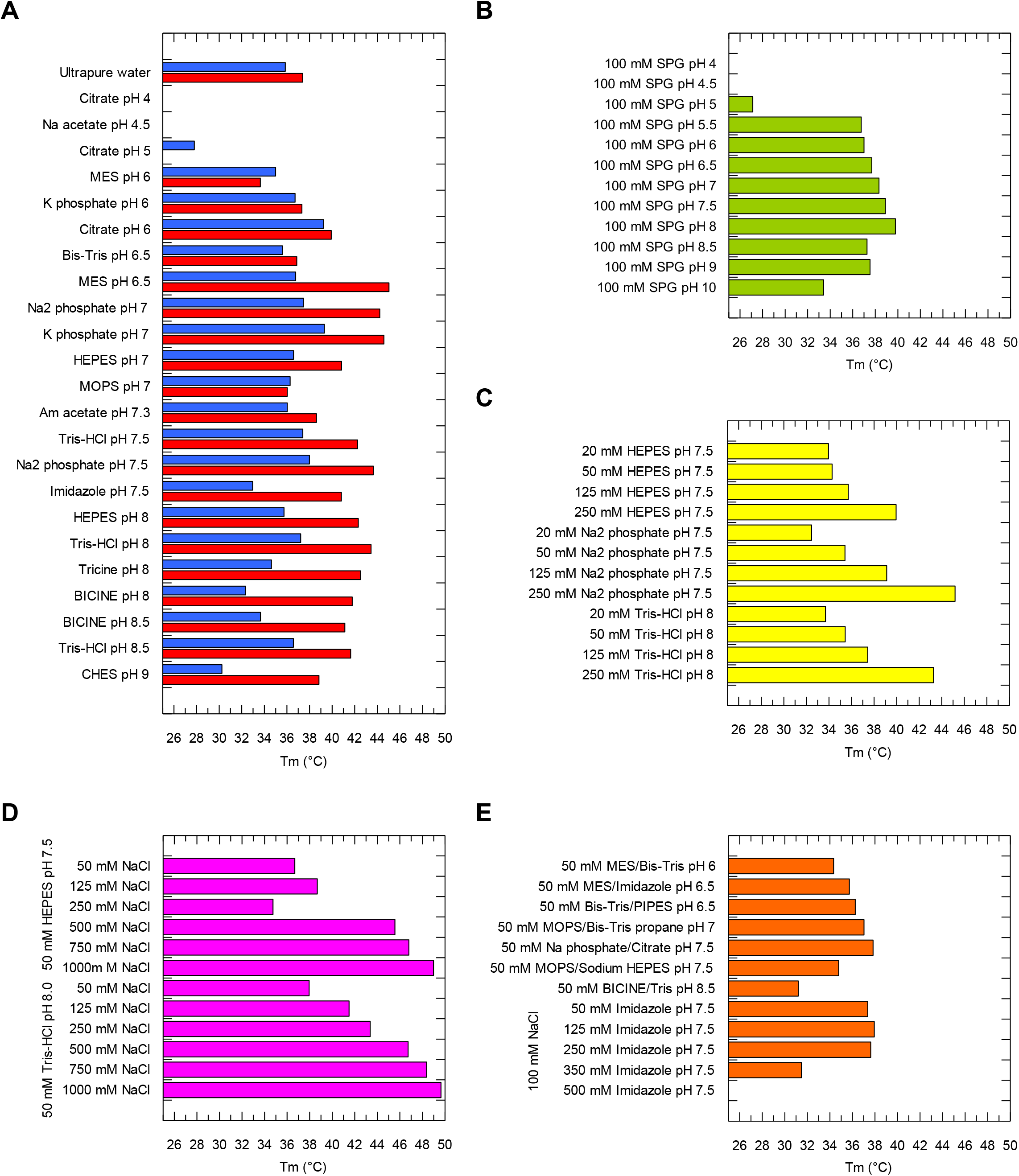
Histograms of TSA T_*m*_ data for all 96 wells. APH(3’)-IIb thermal stability was measured in different conditions using the RUBIC Buffer kit (Molecular Dimensions): (**A**) buffers (100 mM) of different pHs at low (0 mM NaCl, blue bars) or high (250 mM NaCl, red bars) ionic strength, (**B**) composite buffer with extended range of pH deconvoluting pH from buffer effect, (**C**) buffers at different concentrations (20 to 250 mM), (**D**) different ionic strengths by variation of NaCl concentration (50 to 1000 mM) and (**E**) Combination of buffers and effect of imidazole (50 to 500 mM) in presence of 100 mM NaCl.

## Notes

### Competing Interest Statement

The authors have declared no competing interest.

## References

[1] M. Vedadi, F. H. Niesen, A. Allali-Hassani, O. Y. Fedorov, P. J. Finerty, G. A. Wasney, R. Yeung, C. Arrowsmith, L. J. Ball, H. Berglund, R. Hui, B. D. Marsden, P. Nordlund, M. Sundstrom, J. Weigelt, and A. M. Edwards (2006) Chemical screening methods to identify ligands that promote protein stability, protein crystallization, and structure determination. Proc. Natl. Acad. Sci. USA 103, 15835–15840.

[2] L. Reinhard, H. Mayerhofer, A. Geerlof, J. Mueller-Dieckmann, and M. S. Weiss (2013) Optimization of protein buffer cocktails using Thermofluor. Acta Crystallogr. Sect. F Struct. Biol. Cryst. Commun. 69, 209–214.

[3] S. Boivin, S. Kozak, and R. Meijers (2013) Optimization of protein purification and characterization using Thermofluor screens. Protein Expr. Purif. 91, 192–206.

[4] P. H. Lee, X. X. Huang, B. T. Teh, and L. M. Ng (2019) TSA-CRAFT: A Free Software for Automatic and Robust Thermal Shift Assay Data Analysis. SLAS Discov. 24, 606–612.

[5] G. D. Wright (1999) Aminoglycoside-modifying enzymes. Curr. Opin. Microbiol. 2, 499–503.

[6] M. S. Ramirez and M. E. Tolmasky (2010) Aminoglycoside modifying enzymes. Drug Resist. Updat. 13, 151–171.

[7] P. H. Lee (May 2020) TSA-CRAFT: Webserver Implementation. “Hi Victor, Thanks for that you are interested in and implementing our TSA-CRAFT. We have some private issue for maintainance of web version of TSA-CRAFT so that only the standalone version are released at the moment. I am glad that you are so kind and generous to spread TSA-CRAFT for us. Basically, I agree that you/your group to build a public TSA-CRAFT web server. Would you please clearly put the information on your homepage of TSA-CRAFT including the reference of TSA-CRAFT paper, the link of URL for standalone version in source forge and some statement like “the implememtation is under agreement of the authors of TSA-CRAFT”? Regards, Po-Hsien Lee.”.

